# Small molecule inhibition of IRE1α kinase/RNase has anti-fibrotic effects in the lung

**DOI:** 10.1101/398552

**Authors:** Maike Thamsen, Rajarshi Ghosh, Vincent C. Auyeung, Alexis Brumwell, Harold A. Chapman, Bradley J. Backes, Gayani Perara, Dustin J. Maly, Dean Sheppard, Feroz R. Papa

## Abstract

Endoplasmic reticulum stress (ER stress) has been implicated in the pathogenesis of idiopathic pulmonary fibrosis (IPF), a disease of progressive fibrosis and respiratory failure. ER stress activates a signaling pathway called the unfolded protein response (UPR) that either restores homeostasis or promotes apoptosis. The bifunctional kinase/RNase IRE1α is a UPR sensor that promotes apoptosis if ER stress remains high (i.e., a “terminal” UPR). Using multiple small molecule inhibitors against IRE1α, we show that ER stress-induced apoptosis of murine alveolar epithelial cells can be mitigated in vitro. In vivo, we show that bleomycin exposure to murine lungs causes early ER stress to activate IRE1α and the terminal UPR prior to development of pulmonary fibrosis. Small-molecule IRE1α kinase-inhibiting RNase attenuators (KIRAs) that we developed were used to evaluate the importance of IRE1α activation in bleomycin-induced pulmonary fibrosis. One such KIRA—KIRA7—provided systemically to mice at the time of bleomycin exposure decreases terminal UPR signaling and prevents lung fibrosis. Administration of KIRA7 14 days after bleomycin exposure even promoted the reversal of established fibrosis. Finally, we show that KIRA8, a nanomolar-potent, monoselective KIRA compound derived from a completely different scaffold than KIRA7, likewise promoted reversal of established fibrosis. These results demonstrate that IRE1α may be a promising target in pulmonary fibrosis and that kinase inhibitors of IRE1α may eventually be developed into efficacious anti-fibrotic drugs.

## Introduction

The unfolded protein response (UPR) is a conserved signaling pathway that is activated when eukaryotic cells experience protein folding stress in the endoplasmic reticulum (i.e., ER stress). The mammalian UPR is mediated by three sensors of unfolded protein situated in the ER membrane: IRE1α, ATF6, and PERK; of these, IRE1α is the most ancient member as IRE1 orthologs are present in all eukaryotes [1].

Depending on the severity of ER stress, the activity of IRE1α determines cell fate outcomes. When ER stress is remediable, IRE1α homodimerizes in the ER membrane, causing autophosphorylation and selective activation of its C-terminal RNase domain to catalyze the frame-shift splicing of the mRNA encoding the transcription factor XBP1 into its active form (i.e., XBP1s transcription factor) [2–4]. XBP1s in turn promotes homeostasis by upregulating transcription of components of the protein folding machinery including chaperones and post-translational modification enzymes [5,6]. If these adaptive outputs fail to resolve ER stress, IRE1α continues to self-associate into higher-order oligomers on the ER membrane, resulting in kinase hyperphosphorylation and RNase hyperactivation [7,8]. Hyperactivated IRE1α catalyzes endonucleolytic degradation of many messenger RNAs that localize to the ER membrane [7,9]. Hyperactivated IRE1α also cleaves and degrades the precursor of the microRNA miR-17, which in turn relieves the repression of thioredoxin-interacting protein (TXNIP). This pathway culminates in apoptosis and is thus called the “terminal UPR” [10]; terminal UPR signaling underlies several diseases of premature cell loss [11].

Recent studies have implicated the UPR in the pathophysiology of human idiopathic pulmonary fibrosis (IPF), a disease of progressive interstitial fibrosis which leads to severe debilitation and eventually respiratory failure and death [12]. Upregulation of the UPR occurs in both familial IPF and sporadic (non-familial) IPF. In one family, a missense mutation in Surfactant Protein C (SFTPC) causes misfolding of the protein and consequent ER stress in alveolar epithelial cells [13,14], which is recapitulated in a mouse model bearing the same mutation [15]. In types 1 and 4 of the Hermansky-Pudlak syndrome, patients have homozygous loss-of-function at the *HPS1* or *HPS4* loci and develop a devastating pulmonary fibrosis virtually indistinguishable from IPF by early adulthood [16]. *HPS1* or *HPS4* loss-of-function causes accumulation of immature surfactant protein in alveolar epithelial cells and consequently ER stress and apoptosis [17]. Activation of the UPR has also been demonstrated in patients with non-familial (sporadic) IPF [18,19], although the trigger for UPR activation in these patients is unclear. It has been suggested that environmental toxins (cigarette smoke) and infectious agents (e.g. herpesviruses) may increase the secretory workload of alveolar epithelial cells, which may exhaust homeostatic UPR signaling, and ultimately trigger the terminal UPR and cell death [19–21]. In mice, the chemical ER stress inducer tunicamycin is not sufficient to induce fibrosis, but exacerbates fibrosis in the presence of another injury in the form of bleomycin [15]. Administration of the nonspecific chaperones 4-PBA or TUDCA mitigated fibrosis after bleomycin injury [22].

Together, these findings suggest that dampening UPR signaling, perhaps through inhibiting IRE1α, might provide therapeutic benefit in IPF. To these ends, our groups have developed small-molecule compounds that inhibit the IRE1α kinase and thereby allosterically regulate its RNase activity [23–25]. These <u>k</u>inase-inhibiting <u>R</u>Nase <u>a</u>ttenuating (KIRA) compounds dose-dependently reduce IRE1α’s destructive UPR signaling and thereby spare cells experiencing unrelieved ER stress [8,26].

Here we show that ER stress induces apoptosis in a mouse alveolar epithelial cell line and mouse primary type II alveolar epithelial cells, and that inhibiting the IRE1α RNase mitigates apoptosis in vitro. We further show that ER stress and stereotypic terminal UPR signature changes occur in mouse models of bleomycin-induced pulmonary fibrosis, and that administration of a KIRA compound at the time of bleomycin exposure prevents the full fibrosis phenotype. Importantly, we show that KIRA compounds are efficacious even when administered *after* the onset of fibrosis. These studies suggest the possible benefit of the KIRA class of compounds in human IPF.

## Results

### ER stress-induced apoptosis in alveolar epithelial cells depends on IRE1 α activity

Severe or persistent ER stress has been shown to induce apoptosis in diverse cell types [11]. Since chronic epithelial injury is thought to underlie both familial and sporadic IPF, we assessed whether ER stress induces apoptosis in alveolar epithelial cells. The mouse epithelial cell line MLE12 was exposed to tunicamycin, which inhibits N-glycosylation and induces protein misfolding in the endoplasmic reticulum. Tunicamycin induced robust splicing of XBP1 in MLE12 cells, which was dose-dependently inhibited by provision of the covalent IRE1α RNase inhibitor STF-083010 [27] (Fig 1A). Low doses of tunicamycin (30 ng/ml) were sufficient to induce apoptosis in MLE12 cells as measured by annexin V expression (Fig 1B). Tunicamycin-induced apoptosis depended on IRE1α RNase activity, since STF-083010 promoted cell survival. Next, we isolated primary type II alveolar epithelial cells from mouse lungs by dissociation and cell sorting as previously described [28,29]. As in the MLE12 cell lines, tunicamycin induced apoptosis in primary type II alveolar epithelial cells in an IRE1α-dependent manner (Fig 1C).

**Fig 1.**
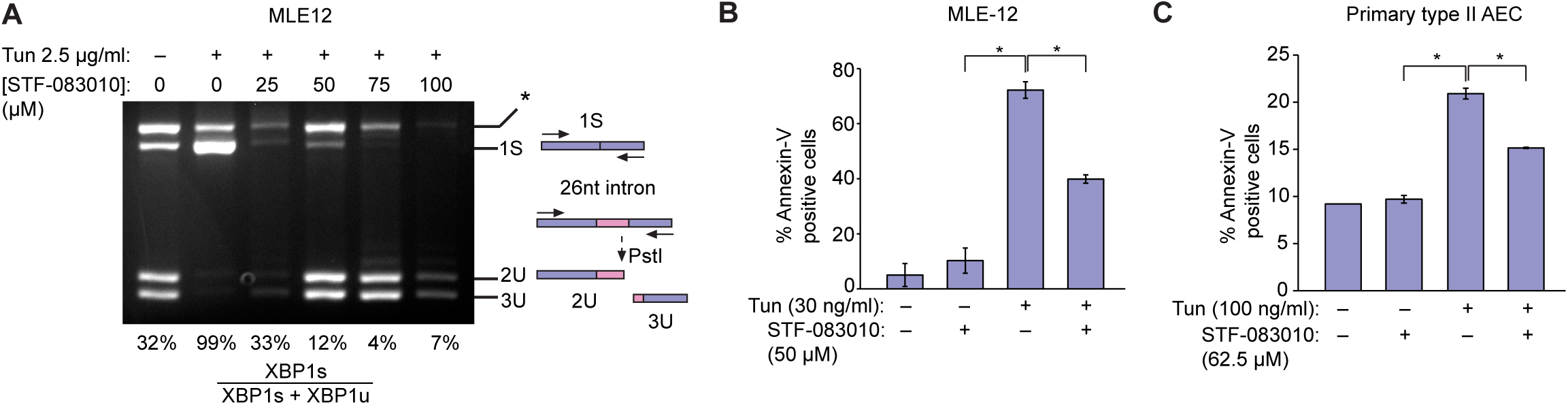
ER stress activates IRE1α to induce apoptosis in alveolar epithelial cells. (A) Ethidium bromide-stained agarose gel of XBP1 cDNA amplicons. Mouse Lung Epithelial (MLE12) cells were exposed to Tunicamycin (Tun) 2.5 μg/ml and indicated concentrations of STF-083010 for 4 hrs. The cDNA amplicon of unspliced XBP1 mRNA is cleaved by a PstI site within a 26 nucleotide intron to give 2U and 3U. IRE1α-mediated cleavage of the intron and re-ligation *in vivo* removes the PstI site to give the 1S (spliced) amplicon. * denotes a spliced/unspliced XBP1 hybrid amplicon. The ratio of spliced over (spliced + unspliced) amplicons—1S/(1S+2U+3U)—is reported as %XBP1 splicing under respective lanes. (B) Percent MLE12 cells staining positive for Annexin V after treatment with Tm (30 ng/ml) and STF-083010 (50 µM) for 72 hrs. (C) Percent primary mouse alveolar epithelial cells (AEC) staining positive for Annexin V after treatment with Tm (100 ng/ml) and STF-083010 (62.5 µM) for 72 hrs. Three independent biological samples were used for Annexin V staining experiments. Data are means +/- SD. P value: * <0.05.

### ER stress precedes fibrosis in the bleomycin model pulmonary fibrosis

To evaluate the role of IRE1α in vivo, we turned to the bleomycin model of pulmonary fibrosis. C57/BL6 mice were exposed to a single dose of intranasal bleomycin (1.5 units/kg), and whole lung RNA was collected at intervals after exposure. Total XBP mRNA levels rise within 2 days of bleomycin exposure (p<0.01), followed by markers of terminal UPR activation: the chaperone BiP (p<0.001), and the transcription factors ATF4 (p<0.01) and CHOP (p<0.05) (Fig 2A). This wave of ER stress and terminal UPR activation peaks at day 8 and remains elevated through day 26 (Fig 2A). This initial, early wave of terminal UPR signaling is followed by overt fibrosis; mRNA levels of collagen 1A1 and fibronectin rise at day 8 and peak on day 14, followed by waning levels over the following two weeks (Fig 2B).

**Fig 2.**
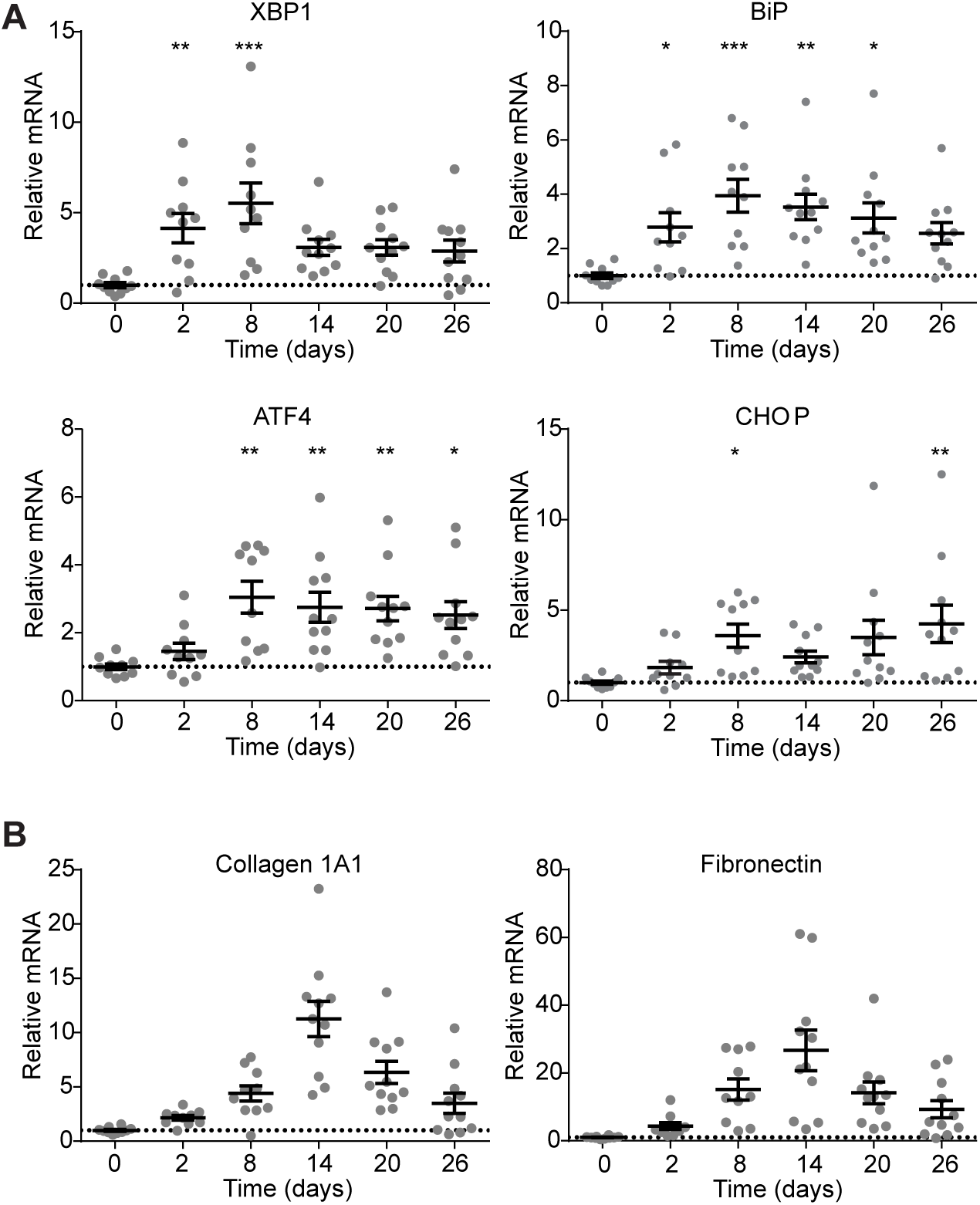
ER stress and terminal UPR signaling precede fibrosis in bleomycin-exposed mice. (A) Quantitative PCR of terminal UPR markers XBP1, BiP, ATF4, and CHOP from mice at 0, 2, 8, 14, 20, or 26 days after a single exposure to bleomycin. Each mouse is represented by a dot, and whiskers denote group mean +/- SEM. (B) Quantitative PCR for collagen 1A1 mRNA and fibronectin mRNA from at 0, 2, 8, 14, 20, or 26 days after a single exposure to bleomycin. Each mouse is represented by a dot, and whiskers denote group mean +/- SEM. P values: *<0.05, **<0.01, ***<0.001.

### KIRA7 modulation of IRE1α prevents bleomycin-induced fibrosis

The timing of ER stress and terminal UPR activation compared to fibrosis onset suggested that administration of an IRE1α inhibitor during the period shortly after bleomycin exposure might mitigate bleomycin-induced fibrosis. The RNase inhibitor STF-083010 is a poor tool compound for in vivo use in part due to its high IC50 of 9.9 µM and because of poor solubility (Selleckchem manufacturer datasheet). Therefore, for in vivo experiments we used KIRA7, an imidazopyrazine compound that binds the IRE1α kinase to allosterically inhibit its RNase activity (Fig 3A). KIRA7, also referred to a Compound 13 [25], is a modification of our previously published compound KIRA6 [8] with comparable potency and pharmacokinetic properties. Mice were exposed to a single dose of intranasal bleomycin (1.5 units/kg), then treated with either KIRA7 (5 mg/kg/day i.p.) or an equivalent volume of vehicle, starting on the day of bleomycin exposure and continuing daily for 14 days. Whole lung protein and RNA were collected on day 14 for analysis (Fig 3B).

**Fig 3.**
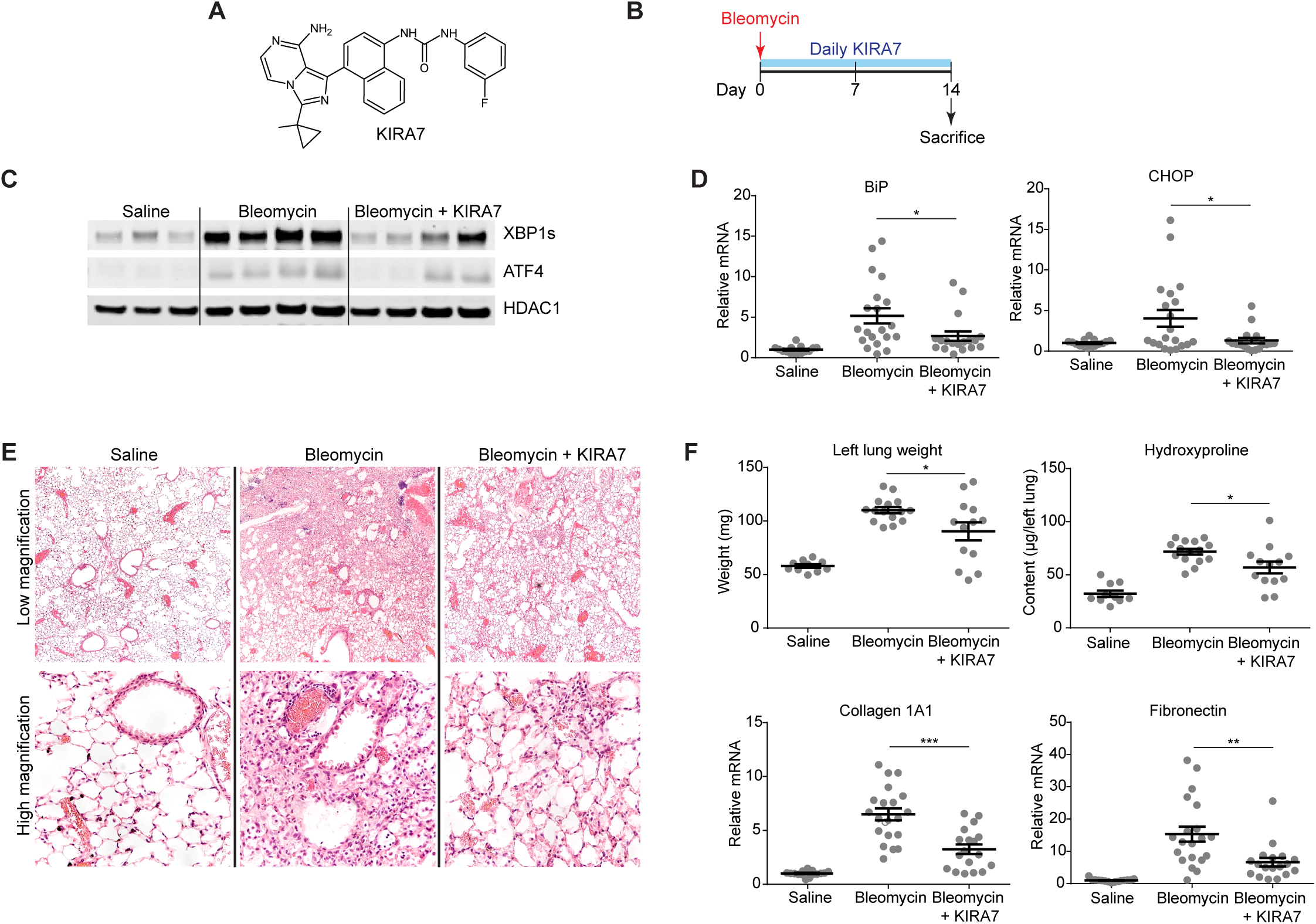
KIRA7 prevents bleomycin-induced fibrosis. (A) Chemical structure of KIRA7. (B) Schematic of the KIRA7 prevention regimen. Mice were exposed to saline or bleomycin once, then treated with KIRA7 or vehicle beginning from the time of bleomycin exposure and daily for two weeks after exposure. (C) Western blot of terminal UPR transcription factors XBP1s and ATF4 from mice treated according to the prevention regimen. (D) Quantitative PCR of terminal UPR markers BiP and CHOP from mice treated according to the prevention regimen. Each mouse is represented by a dot, and whiskers denote group mean +/- SEM. (E) Hematoxylin and eosin stained sections from mice treated according to the prevention regimen, at low magnification (top) and high magnification (bottom). (F) Markers of fibrosis (lung weight, hydroxyproline content, collagen 1A1 mRNA, and fibronectin mRNA) from mice treated according to the prevention regimen. Each mouse is represented by a dot, and whiskers denote group mean +/- SEM. P values: *<0.05, **<0.01, ***<0.001.

Although day 14 is after peak expression of markers of the terminal UPR (Fig 2A), protein levels of spliced XBP1 and ATF4 remain significantly elevated in bleomycin-exposed mice, compared to saline-exposed controls (Fig 3C). Treatment of bleomycin-exposed mice with KIRA7 resulted in decreased spliced XBP1 and ATF4, compared to bleomycin exposed mice treated with vehicle (Fig 3C). Likewise, mRNA levels of BiP and CHOP are significantly elevated after bleomycin exposure, and treatment of bleomycin-exposed mice with KIRA7 decreased these levels (Fig 3D, p<0.05).

These decreases in markers of the terminal UPR were accompanied by significant improvements in bleomycin-induced fibrosis. Histologically, bleomycin exposure induced destruction of alveoli, cell infiltration, and architectural distortion in the lung. By contrast, alveolar architecture was largely preserved in lungs of mice exposed to bleomycin and treated with KIRA7 (Fig 3E). Bleomycin exposure induced fibrosis as measured by lung weight and total hydroxyproline, and both were significantly decreased with KIRA7 treatment (p<0.05). Consistent with these decreases, mRNA levels of collagen 1A1 and fibronectin were both significantly decreased by KIRA7 treatment (p<0.001 and p<0.01, respectively).

### KIRA7 modulation of IRE1 α mitigates established fibrosis

Administration of KIRA7 protects against fibrosis during the initial injury after bleomycin, which is also when ER stress and terminal UPR markers are at their peak (Fig 2A). We wondered whether KIRA7 might even protect against fibrosis when administered later (i.e., in a therapeutic rather than a prophylactic regimen), two weeks after initial injury, when terminal UPR markers are still elevated albeit to a lesser degree (Fig 2A). In addition, early injury in the first week after bleomycin administration is characterized by neutrophilic and lymphocytic inflammatory infiltration, which is not thought to be characteristic of human IPF [30]. The initial alveolitis clears around day 7, followed by fibroblast proliferation and matrix deposition; fibrosis is typically established by day 14 (Fig 2B) [30]. Thus, administration of KIRA7 after day 14 may have greater relevance to human IPF, where fibrosis may progress for years before the onset of clinical symptoms and diagnosis.

As before, mice were exposed to a single dose of intranasal bleomycin (1.5 units/kg). Beginning on day 14, mice were treated with either KIRA7 (5 mg/kg/day i.p.) or an equivalent volume of vehicle; injections were continued daily until day 28. Whole lung protein and RNA were collected on day 28 for analysis (Fig 4A). As expected, bleomycin-exposed mice had increased levels of spliced XBP1, ATF4, and CHOP protein at day 28 (Fig 4B). Treatment of bleomycin-exposed mice was associated with decreases in levels of these proteins (Fig 4B). KIRA7 also blunted bleomycin-induced increases in lung weight (p<0.001) and hydroxyproline (p<0.05) (Fig 4C). At day 28, mRNA levels of collagen 1A1 and fibronectin remain elevated above baseline, albeit at lower levels compared to day 28 (Fig 4C), consistent with the observation that synthesis of collagen and fibronectin mRNA peaks at day 14 and wanes thereafter (Fig 2B). Nonetheless, administration of KIRA7 decreased mRNA levels of collagen 1A1 (p<0.05) and fibronectin (p<0.05).

**Fig 4.**
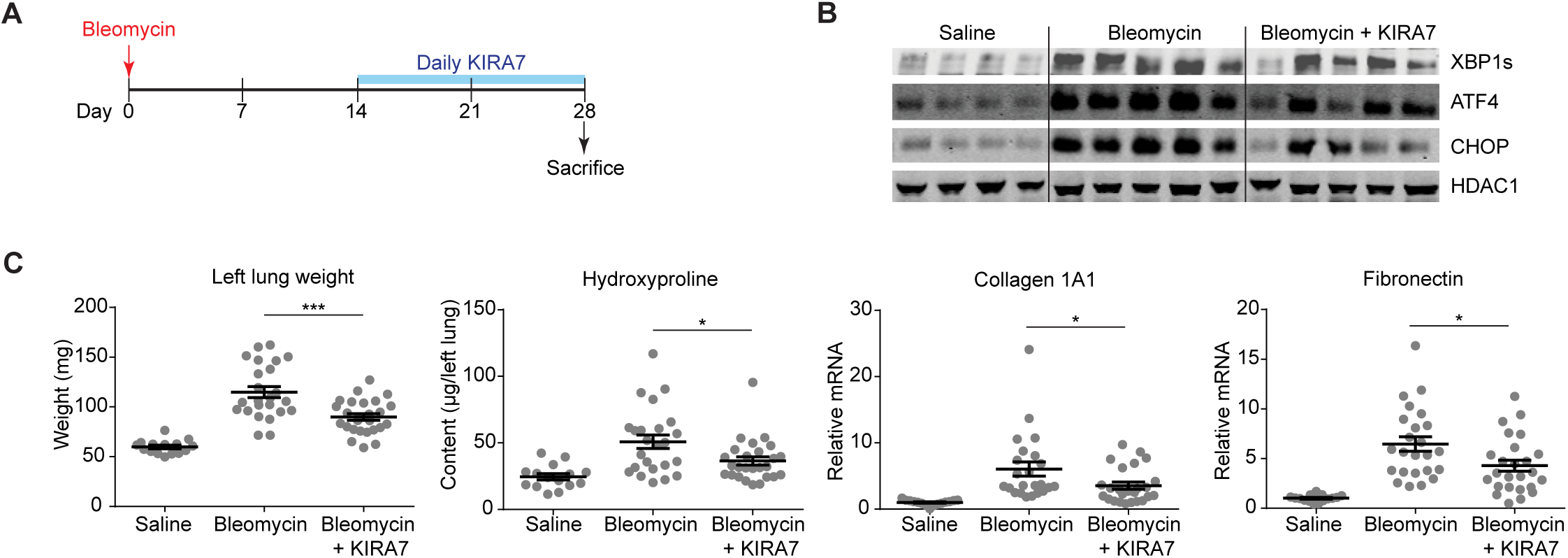
KIRA7 reverses bleomycin-induced fibrosis when given 2 weeks after bleomycin exposure. (A) Schematic of the KIRA7 reversal regimen. Mice were exposed to saline or bleomycin once, then treated with KIRA7 or vehicle beginning two weeks after bleomycin exposure and continuing daily for two additional weeks. (B) Western blot of terminal UPR transcription factors XBP1s, ATF4, and CHOP from mice treated with KIRA7 according to the reversal regimen. (E) Markers of fibrosis (lung weight, hydroxyproline content, collagen 1A1 mRNA, and fibronectin mRNA) from mice exposed exposed to saline or bleomycin, and treated with KIRA7 according to the reversal regimen. Each mouse is represented by a dot, and whiskers denote group mean +/- SEM. P values: *<0.05, **<0.01, ***<0.001.

### KIRA8 modulation of IRE1 α mitigates established fibrosis

A sulfonamide compound was recently described that is structurally unrelated to KIRA7 [31], but possesses the properties of an IRE1α kinase-inactivating RNase attenuator and we therefore call KIRA8 (Fig 5A). It has exceptional selectivity for IRE1α in whole-kinome testing, having little activity even against its closely related paralog IRE1β [26]. KIRA8 is highly potent against the IRE1α kinase (IC50=5.9 nM, [26]), nearly 10-fold more potent than KIRA7 (IC50=46 nM, data not shown). Consistent with this, KIRA8 has higher potency than KIRA7 in inhibiting XBP1 splicing in the alveolar epithelial cell line MLE12 (Fig 5B).

**Fig 5.**
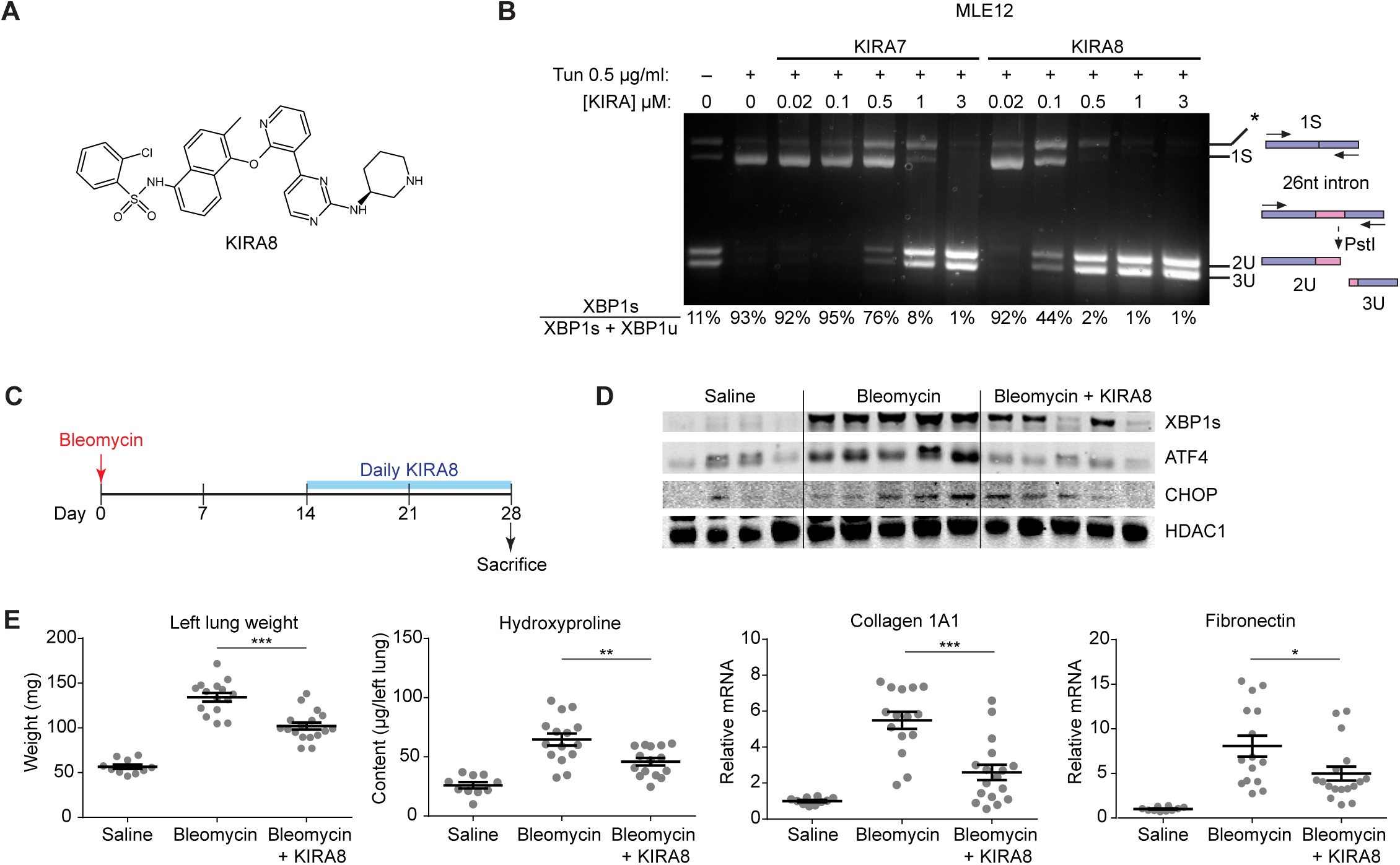
KIRA8 reverses bleomycin-induced fibrosis when given 2 weeks after bleomycin exposure. (A) Chemical structure of KIRA8. (B) EtBr-stained agarose gel of XBP1 cDNA amplicons after induction by treating Mouse Lung Epithelial (MLE12) cells with Tunicamycin (Tun) 0.5 μg/ml and indicated concentrations of KIRA7 or KIRA8 for 8 hrs. The ratio of spliced over (spliced + unspliced) amplicons—1S/(1S+2U+3U)—is reported as % XBP1 splicing and reported under respective lanes. (C) Schematic of the KIRA8 reversal regimen. Mice were exposed to saline or bleomycin once, then treated with KIRA8 or vehicle beginning two weeks after bleomycin exposure and continuing daily for two additional weeks. (D) Western blot of terminal UPR transcription factors XBP1s, ATF4, and CHOP from mice treated with KIRA8 according to the reversal regimen. (E) Markers of fibrosis (lung weight, hydroxyproline content, collagen 1A1 mRNA, and fibronectin mRNA) from mice exposed exposed to saline or bleomycin, and treated with KIRA8 according to the reversal regimen. Each mouse is represented by a dot, and whiskers denote group mean +/- SEM. P values: *<0.05, **<0.01, ***<0.001.

We evaluated the ability of KIRA8 to mitigate established fibrosis. Mice were exposed to a single dose of intranasal bleomycin (1.5 units/kg), and treated with either KIRA8 (50 mg/kg/day i.p.) or an equivalent volume of vehicle starting at day 14 and continuing daily until day 28. Whole lung protein and RNA were collected on day 28 for analysis (Fig 5C). KIRA8 treated mice had lower levels of spliced XBP1, ATF4, and CHOP protein (Fig 5D). As before, KIRA8 treatment blunted bleomycin-induced increases in lung weight (p<0.001) and hydroxyproline (p<0.01), and decreased mRNA expression of collagen 1A1 (p<0.001) and fibronectin (p<0.05) (Fig 5E).

## Discussion

We have shown that ER stress induces apoptosis in a mouse alveolar epithelial cell line and mouse primary type II alveolar epithelial cells, and that inhibiting the IRE1α RNase mitigates apoptosis in vitro. *In vivo*, bleomycin exposure to the mouse lung induces ER stress prior to the onset of fibrosis. Highlighting the importance of ER stress and IRE1α in this model, administration of KIRA7 starting from the time of bleomycin exposure decreased markers of ER stress and prevented fibrosis. Importantly, KIRA7 was efficacious even when administered two weeks after the onset of fibrosis. KIRA8 is a next-generation KIRA compound derived from a completely different scaffold than KIRA7, with nanomolar potency and monoselectivity for the IRE1α kinase. KIRA8 likewise promoted the reversal of established fibrosis.

Several mechanisms may account for the effect of KIRA7 and KIRA8 on bleomycin-induced fibrosis. A prevailing view is that IPF is caused by chronic epithelial injury, which induces fibroblast activation and collagen deposition [32]. In some cases, heritable defects in cargo protein folding or post-translational processing leads to unremediated ER stress, IRE1α activation, terminal UPR signaling, and epithelial cell apoptosis [13,14,16]. Others have proposed that various insults to the alveolar epithelium lead ultimately to terminal UPR signaling [18–21]. In both cases, modulating the activity of IRE1α would be predicted to blunt terminal UPR signaling, promote alveolar epithelial cell survival, and thus mitigate ongoing fibrosis.

Another intriguing possibility is that IRE1α activity may also contribute directly to pathologic fibroblast behavior. For example, fibroblasts derived from patients with systemic sclerosis, IRE1α was required for TGFβ1-induced differentiation into activated myofibroblasts [33]. This finding may help explain why late administration of KIRA7 and KIRA8, two weeks after bleomycin injury, can promote the reversal of established fibrosis (Fig 4 and Fig 5). The possible roles of IRE1α in epithelial cells and fibroblasts are not mutually exclusive, and extensive work outside the scope of this study is needed to elucidate the precise role(s) of IRE1α in the injured lung.

The anti-fibrotic activity of KIRA7 and KIRA8 even when administered late is particularly important when considering potential therapeutic avenues in human disease. In IPF, subclinical fibrosis starts years before patients are symptomatic enough to come to medical attention, and even then diagnosis is often delayed [34,35]. These results suggest that targeting IRE1α may possibly be effective even when fibrosis is advanced. In addition, severe pathological lung fibrosis, evidenced by the “usual interstitial pneumonia” pattern radiographically and histologically, is the final common endpoint to chronic injury in non-IPF interstitial lung diseases, including connective tissue-associated interstitial lung disease, chronic hypersensitivity pneumonitis, and asbestosis [36]. Targeting IRE1α holds therapeutic promise to the extent that ER stress is important to the primary pathology and initial injury in these diseases, as has been suggested in some forms of genetic autoimmune-associated interstitial lung disease [37] and asbestosis [38]. To the extent that targeting IRE1α might mitigate pathological fibroblast activity, targeting IRE1α may be useful even in cases where ER stress is not part of the primary pathology.

In summary, we have shown that intra-airway bleomycin, the most commonly used murine model of pulmonary fibrosis, induces the unfolded protein response and that this response precedes the development of pulmonary fibrosis. Furthermore, two chemically distinct small molecules that inhibit IRE1α kinase and attenuate its RNase activity (KIRAs) are effective in preventing and reversing bleomycin induced pulmonary fibrosis. This work lays the groundwork for developing KIRAs as novel therapeutics for IPF and other interstitial lung diseases characterized by progressive pulmonary fibrosis.

## Materials and methods

### Tissue Culture

Mouse Lung Epithelial (MLE12) cells were obtained from ATCC and grown in HITES media as formulated by ATCC. Tunicamycin (Tun) was purchased from Millipore. STF-083010, KIRA7 and KIRA8 were synthesized in house. Mouse primary AECII cells were isolated as described [39]. Cells were grown in SAGM media (Lonza CC-3118) on fibronectin coated plates.

### XBP-1 mRNA splicing

RNA was isolated from whole cells or mouse tissues and reverse transcribed as above to obtain total cDNA. Then, XBP-1 primers were used to amplify an XBP-1 amplicon spanning the 26 nt intron from the cDNA samples in a regular 3-step PCR. Thermal cycles were: 5 min at 95 °C, 30 cycles of 30 s at 95 °C, 30 s at 60 °C, and 1 min at 72 °C, followed by 72 °C for 15 min, and a 4 °C hold. Mouse XBP1 sense (5’-AGGAAACTGAAAAACAGAGTAGCAGC-3’) and antisense (5’-TCCTTCTGGGTAGACCTCTGG-3’) primers were used. PCR fragments were then digested by PstI, resolved on 3% agarose gels, stained with EtBr and quantified by densitometry using NIH ImageJ.

### Annexin V apoptosis

Annexin V staining was used to quantify apoptosis by flow cytometry. Cells were plated in 12-well plates overnight and then treated with various reagents for indicated time periods. On the day of flow cytometry analysis, cells were trypsinized and washed in PBS and resuspended in Annexin V binding buffer with Annexin-V FITC (BD Pharmingen™). Annexin V stained cells were counted using a Becton Dickinson LSRII flow cytometer.

### Bleomycin-induced pulmonary fibrosis

C57BL6 mice at 12 weeks of age were obtained from Jackson Laboratories. Mice were housed in specific pathogen-free conditions in the Animal Barrier Facility at the University of California, San Francisco. This work was approved by the Institutional Animal Care and Use Committee of the University of California, San Francisco. To induce fibrosis, mice were anesthetized with ketamine and xylazine and exposed to a single dose of intranasal bleomycin (1.5 units/kg). Lungs were harvested at the indicated times. For treatment, KIRA7 or KIRA8 was dissolved in a vehicle consisting of 3% ethanol, 7% Tween-80, and 90% normal saline and injected peritoneally at the indicated dosages and intervals.

Lung hydroxyproline content was quantified by reaction with 4- (Dimethylamino)benzaldehyde reaction and colorimetry (Sigma). Messenger RNA levels were measured by reverse transcription and quantitative PCR as described below.

### RNA isolation and quantitative real-time PCR (qPCR)

A TissueLyser II (Qiagen) was used to homogenize mouse lungs for RNA isolation. RNA was isolated from lung homogenates or cultured cells using either Qiagen RNeasy kits or Trizol (Invitrogen). For cDNA synthesis, 1 μg total RNA was reverse transcribed using the QuantiTect Reverse Transcription Kit (Qiagen). For qPCR, we used SYBR green (Qiagen) and StepOnePlus Real-Time PCR System (Applied Biosystems). Thermal cycles were: 5 min at 95 °C, 40 cycles of 15 s at 95 °C, 30 s at 60 °C. Gene expression levels were normalized to 18S rRNA. Primers used for qPCR were as follows:

Human/Mouse 18S rRNA: 5’-GTAACCCGTTGAACCCCATT-3’ and 5’- CCATCCAATCGGTAGTAGCG-3’

Mouse XBP1: 5’-CCGTGAGTTTTCTCCCGTAA-3’ and 5’-AGAAAGAAAGCCCGGATGAG-3’

Mouse BiP: 5’-TCAGCATCAAGCAAGGATTG-3’ and 5’-AAGCCGTGGAGAAGATCTGA-3’

Mouse ATF4: 5’-GCAAGGAGGATGCCTTTTC-3’ and 5’-GTTTCCAGGTCATCCATTCG-3’

Mouse CHOP: 5’-CACATCCCAAAGCCCTCGCTCTC-3’ and 5’- TCATGCTTGGTGCAGGCTGACCAT-3’

Mouse Collagen 1A1: 5’-CCTGGTAAAGATGGTGCC-3’ and 5’- CACCAGGTTCACCTTTCGCACC-3’

Mouse Fibronectin: 5’-ACAGAAATGACCATTGAAGG-3’ and 5’-TGTCTGGAGAAAGGTTGATT- 3’

### Western blot

Whole lungs were homogenized using a TissueLyzer II (Qiagen). Nuclear and cytoplasmic fractions were isolated using the NE-PER extraction kit (Thermo Fisher). Western blots were performed using 4%-12% Bis-Tris precast gels (Invitrogen) using MOPS buffer, then transferred onto nitrocellulose membranes. Antibody binding was detected using conjugated secondary antibodies (Li-Cor) on the Li-Cor Odyssey scanner. Antibodies used for Western blot were as follows: XBP1 (Biolegend 9D11A43), HDAC1 (Cell Signaling Technologies 5356), ATF4 (Sigma WH0000468M1), TXNIP (MBL International K0205-3), and CHOP (Cell Signaling Technologies 2895).

